# Is there a role for the RNA-binding protein LARP1 in β-cells?

**DOI:** 10.1101/2020.09.03.281832

**Authors:** Joao Pedro Werneck-de-Castro, Flavia Leticia Martins Peçanha, Diego Silvestre, Ernesto Bernal-Mizrachi

## Abstract

Mechanistic target of rapamycin complex 1 (mTORC1) is a cellular rheostat linking nutrient availability and growth factor to cellular protein translation. In pancreatic insulin secreting β-cells, mTORC1 deficiency or chronic hyperactivation leads to diabetes. mTORC1 complexes with La-related protein 1 (LARP1) to specifically regulate the expression of 5’ terminal oligopyrimidine tract (5’TOP) mRNAs which encode proteins of the translation machinery and ribosome biogenesis. We aimed to investigate the role played by LARP1 in β-cells *in vivo*. Here we show that LARP1 is the most expressed LARP in mouse islets and human β-cells, being 2-4-fold more abundant than LARP1B, a member of the family that also interacts with mTORC1. Interestingly, β-cells from diabetic patients have higher LARP1 and LARP1B expression suggesting greater protein translation. These studies led us to generate a conditional LARP1 knockout mouse in β-cells (*β-Larp1KO* mice). These mice exhibit normal levels of all LARP family members including *Larp1B, Larp4, Larp6* and *Larp7*. We did not observe any difference between control and *β-Larp1KO* male mice in body weight gain, glucose levels and glucose tolerance at 8, 14 and 44 weeks of age. Female *β-Larp1KO* mice also performed normally during the glucose tolerance test. We then challenged the *β-Larp1KO* mice with high fat (HFD) or high branched-chain amino acid (BCAA) diets. During the course of 8 weeks in HFD, *β-Larp1KO* and control mice had similar weight gain and did not show alterations in glucose homeostasis compared to control littermates. BCAA did not impair glucose metabolism up to 8 weeks of diet challenge. However, glucose tolerance was slightly impaired in the *β-Larp1KO* mice at 16 weeks under BCAA diet. In conclusion, LARP1 is the most abundant LARP in mouse islets and human β-cells and it is upregulated in diabetic subjects. While the lack of LARP1 specifically in β-cells did not alter glucose homeostasis in basal conditions, long-term high branched-chain amino acid feeding could impair glucose tolerance.

## INTRODUCTION

Insulin secreting β-cell failure is a hallmark of diabetes (Alejandro, Gregg et al. 2015). Although β-cells are competent in adapting to insulin resistance by secreting more insulin and increasing in mass to maintain glucose homeostasis, the high metabolic demand will eventually progress to β-cell exhaustion and a fraction of obese patients will develop diabetes (Chang-Chen, Mullur et al. 2008, Alejandro, Gregg et al. 2015). Mechanistic target of rapamycin complex 1 (mTORC1) is a cellular rheostat linking nutrient availability and growth factor signaling to cell metabolism. We have shown previously that mTORC1 is essential to β-cell function and mass (Alejandro, Bozadjieva et al. 2017, Blandino-Rosano, Barbaresso et al. 2017). Lack of mTORC1 activity specifically in β-cells impairs proliferation and survival (Blandino-Rosano, Barbaresso et al. 2017). mTORC1 also regulates insulin processing (Blandino-Rosano, Scheys et al. 2016, Alejandro, Bozadjieva et al. 2017, Blandino-Rosano, Barbaresso et al. 2017) as well as β-cell maturation (Ni, Gu et al. 2017, Jaafar, Tran et al. 2019, Helman, Cangelosi et al. 2020). On the other hand, chronic hyperactivation of mTORC1 in β-cells culminates into β-cell dysfunction and diabetes (Bartolome, Kimura-Koyanagi et al. 2014, Ardestani, Lupse et al. 2018). These findings underscore the importance of mTORC1 signaling in β-cells.

Protein translation depends on an orchestrated assembling of proteins participating in translation initiation, ribosomal recruitment and protein elongation. mTORC1 controls cell size, proliferation, ribosomal biogenesis, protein translation and autophagy by modulating eIF4E-binding proteins (4E-BP1, 2 and 3) and ribosomal protein S6 kinases (S6K1 and 2) and ULK among others (Shimobayashi and Hall 2014, Efeyan, Comb et al. 2015). It also enhances cellular protein synthesis capacity by inducing translation of certain 5’ terminal oligopyrimidine tract (5’TOP) mRNAs which encode proteins of the translation machinery and ribosome biogenesis (Jefferies, Fumagalli et al. 1997, Thoreen, Chantranupong et al. 2012, Meyuhas and Kahan 2015, Hinnebusch, Ivanov et al. 2016). mTORC1 inhibition by rapamycin represses TOP mRNA translation (Jefferies, Reinhard et al. 1994, Terada, Patel et al. 1994, Jefferies, Fumagalli et al. 1997) and the 4EBP proteins have been proposed as suppressors of TOP mRNAs translation (Thoreen, Chantranupong et al. 2012). Recently, a downstream target of mTORC1, the La-related protein 1 (LARP1), also known as RNA-binding protein LARP1, has been described as part of the protein complex regulating the 5’-TOP mRNA translation (Tcherkezian, Cargnello et al. 2014, Deragon and Bousquet-Antonelli 2015, Fonseca, Zakaria et al. 2015, Mura, Hopkins et al. 2015, Hong, Freeberg et al. 2017, Lahr, Fonseca et al. 2017). The LARP family consists of six members: LARP1, 2 (1B), 4, 5 (4B), 6, and 7 (Bousquet-Antonelli and Deragon 2009, Hong, Freeberg et al. 2017). They all contain the RNA-binding domain but only LARP1 and LARP1B present the DM15 motif and interact with mTORC1 (Bousquet-Antonelli and Deragon 2009, Hong, Freeberg et al. 2017).

The role of LARP1 in protein synthesis and mTORC1-mediated TOP mRNA translation is controversial. Studies in HeLa and HEK293 cells have demonstrated that LARP1 positively regulates protein synthesis (Burrows, Abd Latip et al. 2010, Aoki, Adachi et al. 2013, Tcherkezian, Cargnello et al. 2014). Knockdown of LARP1 impaired cell division and migration, and induces cell apoptosis as well as a 15% drop in overall protein synthesis accompanied by hypophosphorylated 4E-BP1 levels (Burrows, Abd Latip et al. 2010), indicating participation in cap-dependent translation. LARP1 depletion was associated with a decreased TOP mRNAs translation (Tcherkezian, Cargnello et al. 2014). In contrast, also using HEK293T and HeLa cells, Fonseca et al. concluded that LARP1 functions as an important repressor of TOP mRNA translation downstream of mTORC1 (Fonseca, Zakaria et al. 2015). They showed that LARP1 interacts with TOP mRNAs in an mTORC1-dependent manner and competes with the eIF4G for TOP mRNA binding. Reducing LARP1 protein levels by siRNA attenuates the inhibitory effect of rapamycin and Torin1 on TOP mRNA translation (Fonseca, Zakaria et al. 2015). That LARP1 directly binds the cap and adjacent 5’TOP motif of TOP mRNAs impeding access of eIF4E and eIF4F assembly was confirmed later (Lahr, Fonseca et al. 2017). Phosphorylation of LARP1 by mTORC1 and Akt/S6K1 dissociates it from 5’UTRs and relieves its inhibitory activity (Hong, Freeberg et al. 2017). Concomitantly, phosphorylated LARP1 scaffolds mTORC1 on the 3’UTRs to facilitate mTORC1-dependent induction of translation initiation. Although LARP1 has inhibitory effects on TOP mRNA translation, LARP1 loss of function causes inefficient translation elongation (Hong, Freeberg et al. 2017). The *in vivo* role of LARP proteins has been limited by the lack of animal models to study the tissue specific importance of this molecule.

We document herein that LARP1 is the most abundant of the family in human β-cells and mouse islets by our own experiments and by assessing public RNA expression databases. Interestingly, diabetes increases LARP1 and LARP1B in human β-cells, suggesting higher protein translation capacity and consistence with greater mTORC1 activity in diabetes. To study the role of LARP1 in β-cells, we developed mice with conditional inactivation in pancreatic β-cells. We found that *Larp1* gene disruption in mouse β-cells is dispensable for glucose homeostasis during normal conditions or after administration of high fat. Challenging the *β-Larp1KO* mice with high branched-chain amino acid diets only impaired glucose tolerance after 16 weeks. We conclude that although LARP1 is highly expressed in β-cells and is up-regulated in diabetogenic conditions, yet it is not essential for glucose homeostasis in normal conditions. LARP1 could play a role in longterm high branched-chain amino acid feeding.

## METHODS

### Human β-cells gene expression database

In order to determine LARP1 family gene expression in human β-cells, we assessed publicly available databases (Blodgett, Nowosielska et al. 2015, Segerstolpe, Palasantza et al. 2016). LARP family gene expression in human fetal and adult β-cells were calculated from Blodgett et al. (Blodgett, Nowosielska et al. 2015). In addition, single-cell transcriptome of healthy and type 2 diabetic subjects’ database by Segerstolpe et al. (Segerstolpe, Palasantza et al. 2016) was used to assess the expression of LARP family members in β-cells.

### Mice

All the procedures were approved by University of Miami IACUC committee (IACUC protocol #18-168). To generate the *floxed-Larp1* mouse, embryonic stem cells containing the floxed *Larp1* construct (Fig 3A) were obtained from the International Mouse Phenotype Consortium (mousephenotype.org/data/genes/MGI:1890165; Larp1^tm1a(EUCOMM)Hmgu^) and injected into blastocyst to generate chimeric mice by the University of Michigan Transgenic Animal Model core (Alejandro, Bozadjieva et al. 2017). After identifying germline transmission, founder lines were selected and bred into *C57BL/6J* background. In order to obtain the LARP1 knockout mice specifically in β-cells (*β-Larp1KO*), *floxed-Larp1* mice were crossed with mice expressing cre-recombinase under the rat insulin promoter activity (*RIP-Cre^Herr^* mice (Alejandro, Bozadjieva et al. 2017, Blandino-Rosano, Barbaresso et al. 2017). We also disrupted *Larp1* gene by crossing the *Floxed-Larp1* mice with the *Ins1-cre* mouse (B6(Cg)-Ins1tm1.1(cre)Thor/J; Jackson’s lab stock no:026801).

### Metabolic studies

Blood glucose levels were determined from blood obtained from the tail vein using ACCU-CHEK II glucometer (Roche). Glucose tolerance test was performed in 6 h fasted animals by injecting glucose intraperitoneally (2 g/kg).

### Diets

High Fat Diet (HFD; cat no D12492; 20 % of carbohydrate, 20 % protein and 60 % fat; Research Diets, New Brunswick, NJ) and Branched-chain amino acid enriched diet (BCAA; cat no D07010503; 67 % of carbohydrate, 23 % protein and 10% fat Research Diets). The BCAA has 150% more leucine, isoleucine and valine concentrations. Control standard diet contains 55 % of carbohydrate, 23 % protein and 22 % fat (ENVIGO).

### Pancreatic islets isolation and quantitative real-time PCR

Islets were isolated by collagenase digestion method as detailed recently (Werneck-de-Castro, Blandino-Rosano et al. 2020). After overnight recover in non-treated plastic petri dishes (VWR) containing RPMI 1640 (Corning cellgro) supplemented with 10% FBS, 1% Penicillin and Streptomicin and 5.5 mM glucose, total RNA was isolated using RNeasy (Qiagen) from 80-100 islets. cDNA was synthesized from 0.5 ug of total RNA using random hexamers and was reverse transcribed using Superscript II (High Capacity cDNA reverse transcription kit; Applied biosystems). Real-time PCR was performed on an ABI 7000 sequence detection system using POWER SYBR-Green PCR Master MIX (Applied Biosystems). Primers were purchased from IDT Technologies and sequences are shown in Table 1.

**Table 1.**
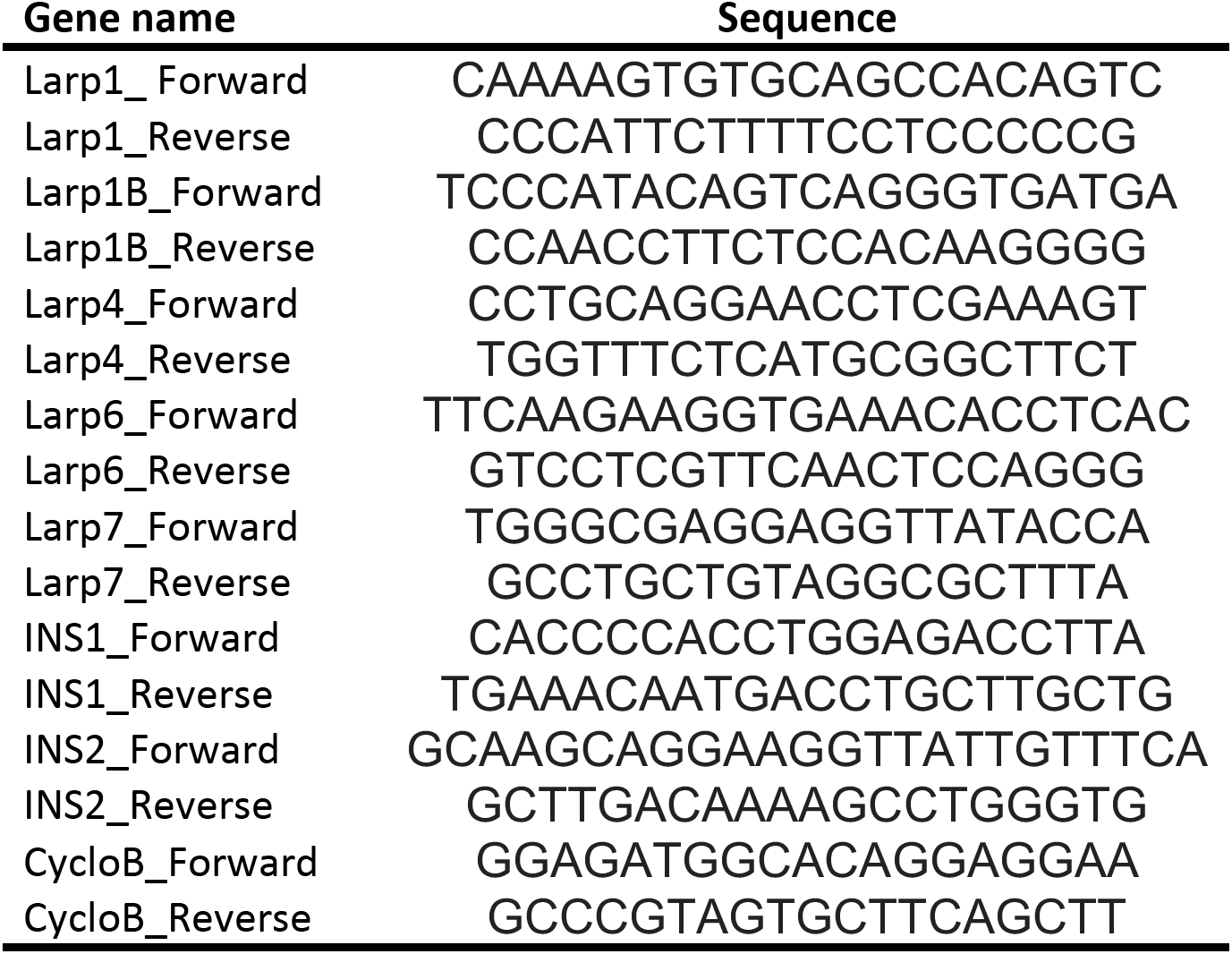
qRT-PCR primer sequences.

### Statistics

Data are presented as mean ± sem. Student t test was employed to assess statistical difference between means of two groups in one time point, e.g. control vs diabetes (Figure 2C) and control vs *β-Larp1KO* mice (Fig 3B). One-way analysis of variance (*ANOVA*) followed by Dunnet’s posthoc test was performed to compare LARP family gene expression to the LARP1 mRNA levels (Fig 2A and B). Two-way *ANOVA* followed by Tukey’s posthoc test was used to identify differences between control vs *β-Larp1KO* mice over time, e.g. body weight gain (Fig 4A, 6A and 7A) and glucose levels during intraperitoneal and oral glucose tolerance test. Results were considered statistically significant when the p value was less than 0.05.

## RESULTS

### LARP1 is the most expressed LARP in human and mouse pancreatic islets and β-cells

The RNA-binding La-related protein (LARP) family consists of six members namely LARP1, 2 (1B), 4, 5 (4B), 6, and 7 (Bousquet-Antonelli and Deragon 2009, Hong, Freeberg et al. 2017). LARP1 and LARP1B share a common DM15 motif and are mTORC1 targets. We first measured the mRNA levels of the LARP family in isolated mouse islets (Fig 1A). Remarkably, LARP1 is the most expressed LARP in mouse islets, being 4-5-fold higher than LARP1B, LARP4 and LARP7 (Fig 1A). LARP6 is barely detectable, beeing only 5% of LARP1 expression (Fig 1A).

**Figure 1.**
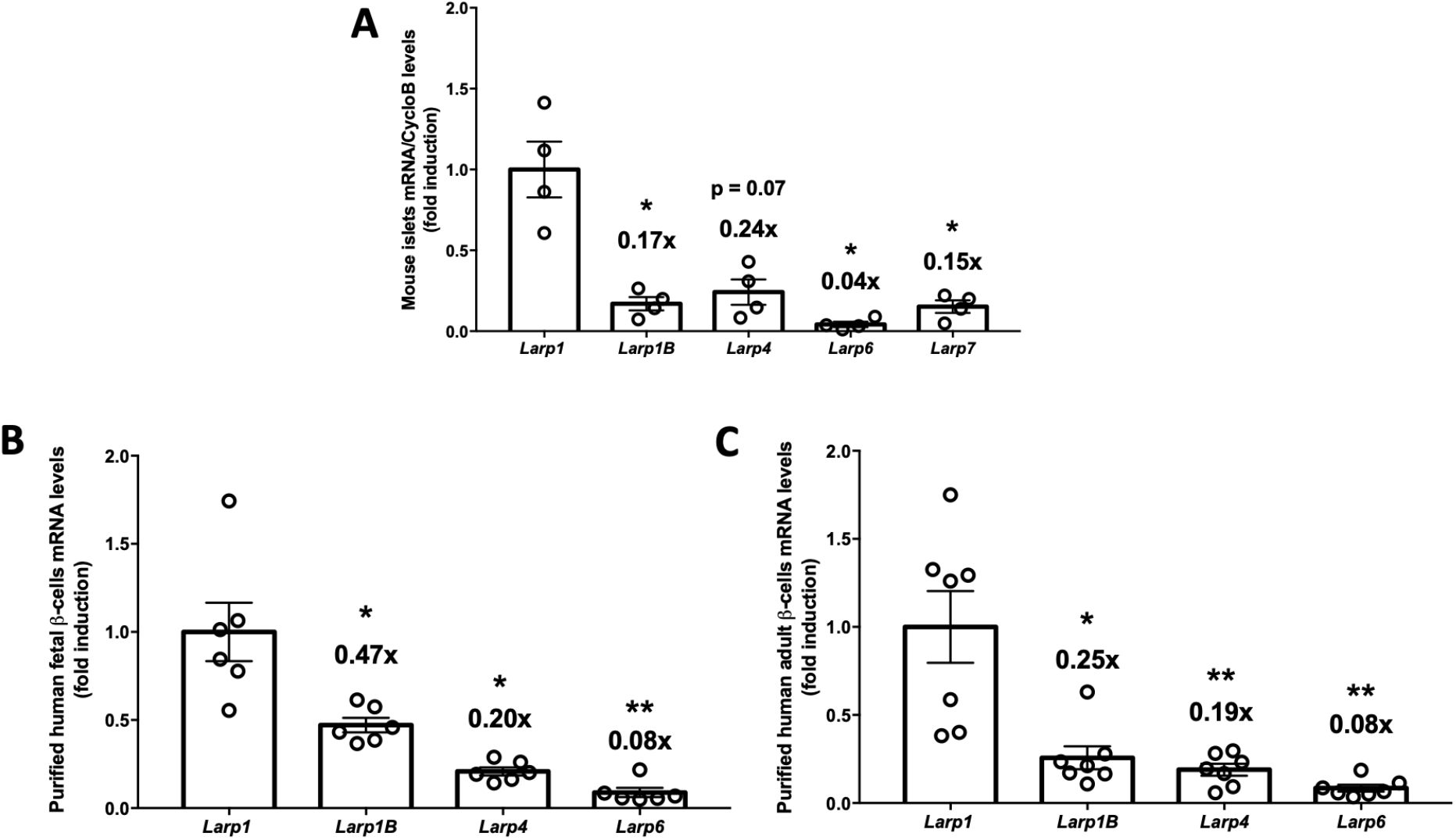
LARP1 is the most expressed LARP in mouse islets and human β-cells. **(A)** LARP family mRNA levels in isolated mouse islets, **(B)** LARP family gene expression in human fetal β-cells and **(C)** Same as in B except that adult human β-cells were used * p < 0.05, ** p < 0.01 and *** p < 0.001 compared to LARP1 mRNA levels assessed by one-way analysis of variance (*ANOVA*) followed by Dunnet’s posthoc test. Numbers on top of the bars denote fold reduction. Figures B and C are analysis of RNA sequencing publicly available database by Blodgett et al. (Blodgett, Nowosielska et al. 2015).

Assessment of publicly available mRNA transcription databases for human β-cells (Blodgett, Nowosielska et al. 2015, Segerstolpe, Palasantza et al. 2016) shows that fetal and adult purified β-cells present higher levels of LARP1 mRNA levels (~2-fold) compared to its paralog gene LARP1B (Fig 1B and C; (Blodgett, Nowosielska et al. 2015)). Single-cell transcriptome profiling of human β-cells in healthy subjects and type 2 diabetes (T2D) further corroborates that LARP1 is the most abundant LARP in β-cells (Fig 2A) with similar pattern seen in T2D (Fig 2B). Interestingly, diabetes increases LARP1 (~30%) and LARP1B (100%) (Fig 2C), suggesting higher protein translation capacity, consistent with greater mTORC1 activity in diabetes (Ardestani, Lupse et al. 2018).

**Figure 2.**
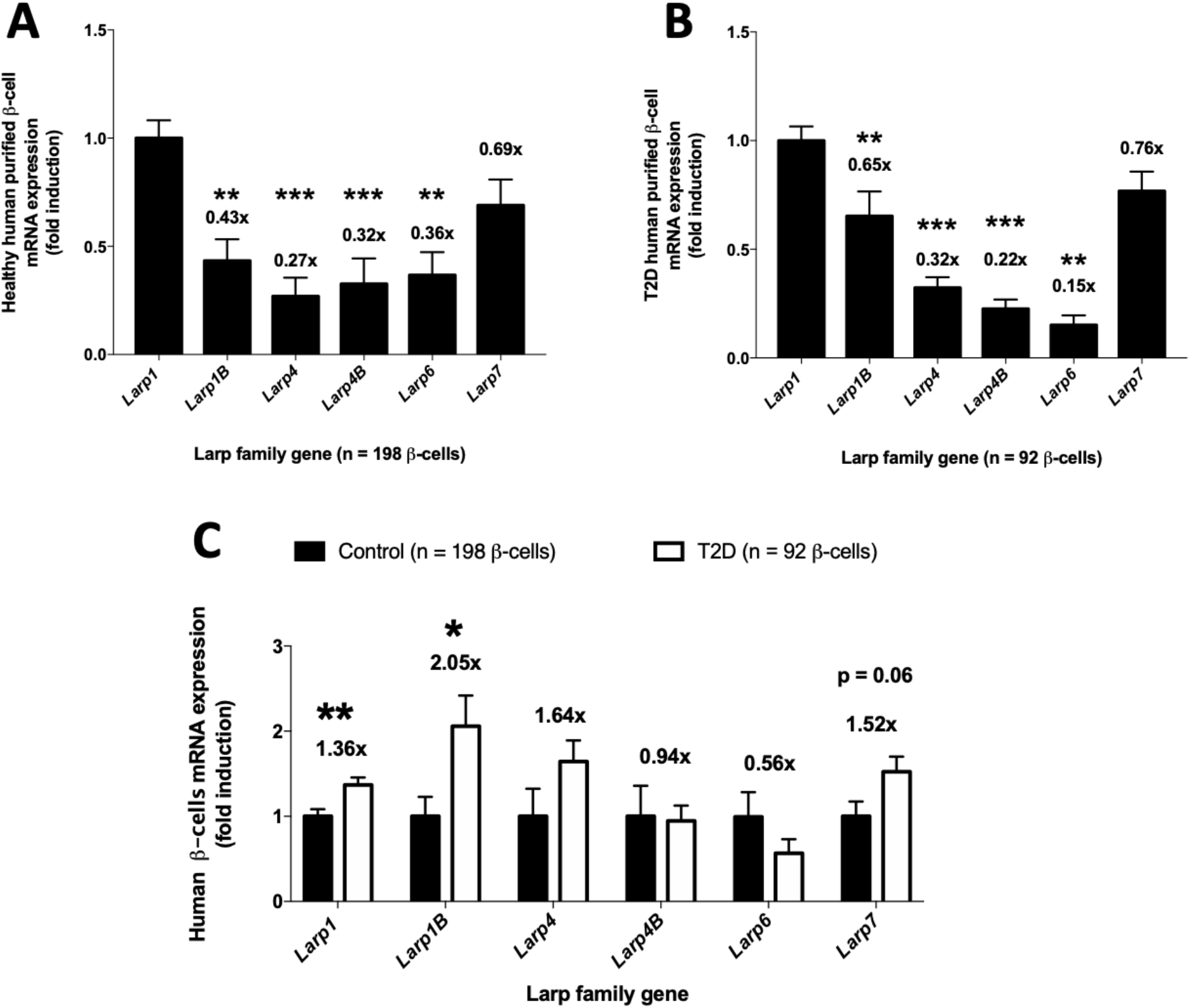
Diabetes increases Larp1 and Larp1B expression in human β-cells. **(A)** LARP family mRNA levels in human β-cells of healthy subjects. **(B)** Same as in A except that human β-cells from type 2 diabetic patients were used. ** p < 0.01 and *** p < 0.001 compared to LARP1 mRNA levels assessed by one-way analysis of variance (*ANOVA*) followed by Dunnet’s posthoc test. **(C)** Fold induction of LARP family mRNA levels in β-cells from type 2 diabetes compared to healthy control subjects shown in A and B. * p < 0.05 and ** p < 0.01 compared to healthy subjects assessed by Student’s T test. Numbers on top of the bars denote fold induction. All data were obtained by analysis of single-cell transcriptome available by Segerstolpe et al. (Segerstolpe, Palasantza et al. 2016)

### β-Larp1KO mice: in vivo model to assess LARP1 function

Whereas the role played by mTORC1 in β- and a-cells was recently revealed (Alejandro, Bozadjieva et al. 2017, Blandino-Rosano, Barbaresso et al. 2017, Bozadjieva, Blandino-Rosano et al. 2017, Ni, Gu et al. 2017, Sinagoga, Stone et al. 2017), the role of the mTORC1 target LARP1 on pancreatic β-cells has never been studied. mTORC1 complexes and phosphorylates LARP1, but whether LARP1 inhibits or stimulates mTORC1-mediated protein translation of TOP mRNAs is still under debate (Burrows, Abd Latip et al. 2010, Aoki, Adachi et al. 2013, Tcherkezian, Cargnello et al. 2014, Fonseca, Zakaria et al. 2015, Mura, Hopkins et al. 2015, Hong, Freeberg et al. 2017, Lahr, Fonseca et al. 2017). Published work has characterized the role of LARP1 using *in vitro* models. Therefore, we decided to generate a LARP1 deficient mouse specifically in β-cells to evaluate LARP1 function in the context of β-cell biology. We generated a*floxed-LARP1* mouse with the lox p sequence flanking exon 4 (Figure 3A). Then, *floxed-LARP1* mice were crossed initially with mice expressing Cre recombinase under the control of the rat insulin promoter to generate the *β-Larp1KO* mouse. LARP1 mRNA levels decreased 80% in isolated islets of the *β-Larp1KO* compared to littermate control mice (Fig 3B). The remaining expression is probably from non-*β*-cells (e.g. a and δ cells) and acinar contaminants. The expected recombination in *β*-cells with the RIP-cre mouse is about 90-95% (Blandino-Rosano, Barbaresso et al. 2017), therefore, the residual expression can also be explained by the non-recombined *β*-cells. LARP1 and LARP1B are very similar in structure sharing the DM15 region, known as LARP1 motif (Stavraka and Blagden 2015, Hong, Freeberg et al. 2017). Therefore, we measured LARP1B and other LARP family members mRNA expression in isolated islet. The deficiency is specific to LARP1, since the *β-Larp1KO* mice have normal expression of LARP1B, LARP4 and LARP7. LARP6 mRNA levels tended to be increased in the *β-Larp1KO* mice but did not reach statistical significance (Fig 3B).

**Figure 3.**
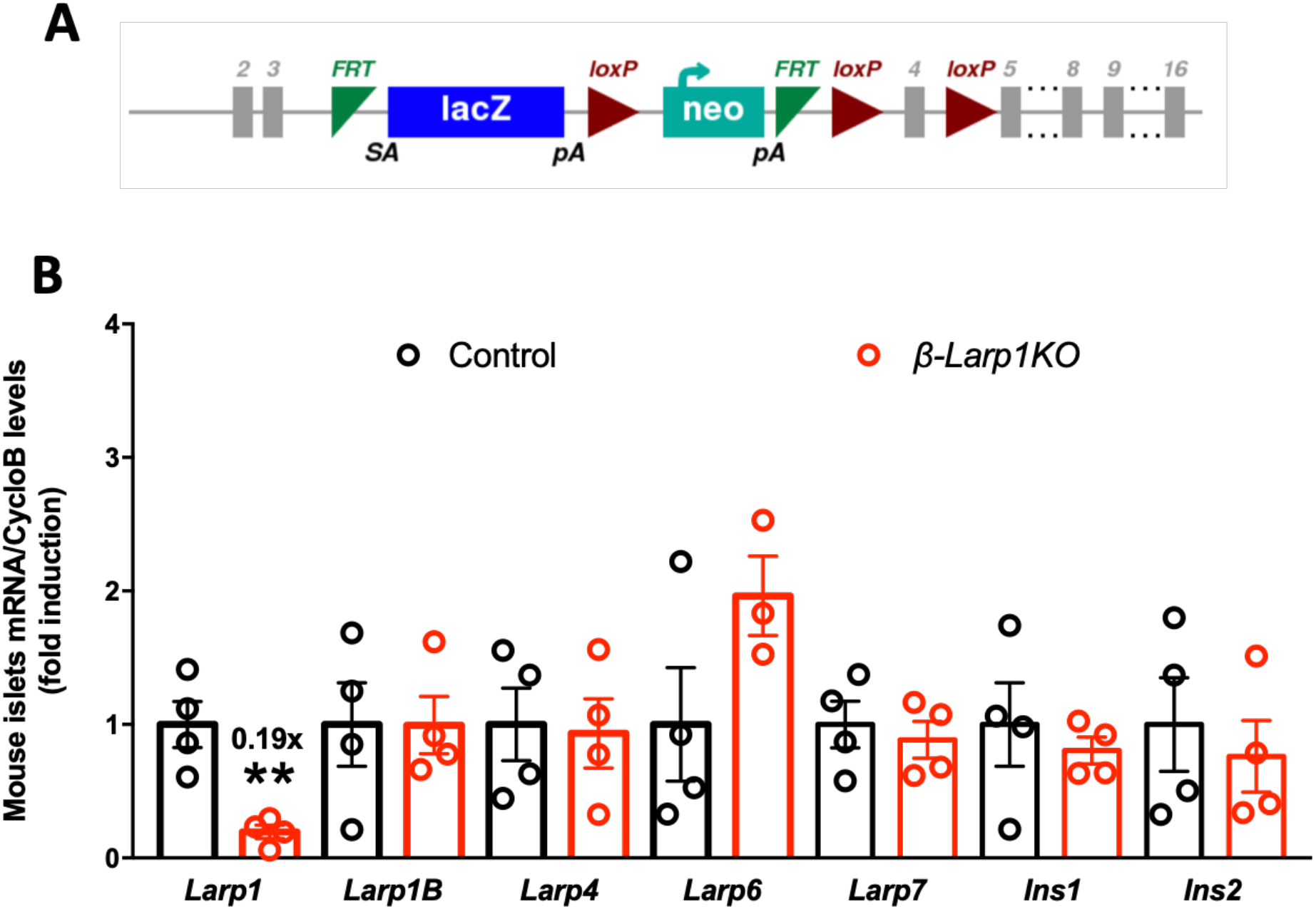
Generation of conditional knockout mice in β-cells (β-Larp1KO). **(A)** *Larp1* gene construct used to generate the *floxed-Larp1* mouse. Cre-recombinase target Loxp sequences were inserted flanking exon 4. **(B)** LARP family and insulin mRNA levels in isolated islets from control and *β-Larp1KO* mice. ** p < 0.01 compared to control mice assessed by Student’s T test. Number on top of *Larp1* bar denote fold induction.

### β-Larp1KO mice grow and age normally and exhibit normal glucose homeostasis

We followed up glucose metabolism in *β-Larp1KO* male mice at different ages (Fig 4). Body weight, fed and fasting glycemia, and tolerance to intraperitoneal glucose load were similar in the first cohort of animals at 4 and 8 weeks of age (Fig 4A-C). Using a second cohort of *β-Larp1KO* male mice, no difference in body weight, fed and fasting glycemia and glucose tolerance test was observed at 14 weeks of age (Fig 4D-F). In addition, there was no difference in glucose tolerance in *β-Larp1KO* female mice (Fig 4G). We also crossed the *floxed-LARP1* mouse with *Ins1-Cre* knock-in mice (Ins1-cre) and found no difference in glucose tolerance (Fig 4H) despite the 80% reduction in Larp1 expression by islets (Fig 4I). To investigate the effects of aging and the lack of LARP1 in *β*-cells, glucose tolerance test was performed in a third cohort of mice aged until almost a year (Fig 5). Aged *β-Larp1KO* male mice performed similarly to control mice in the glucose tolerance test and weighted equally at 44 weeks of age (Fig 5A and B).

**Figure 4.**
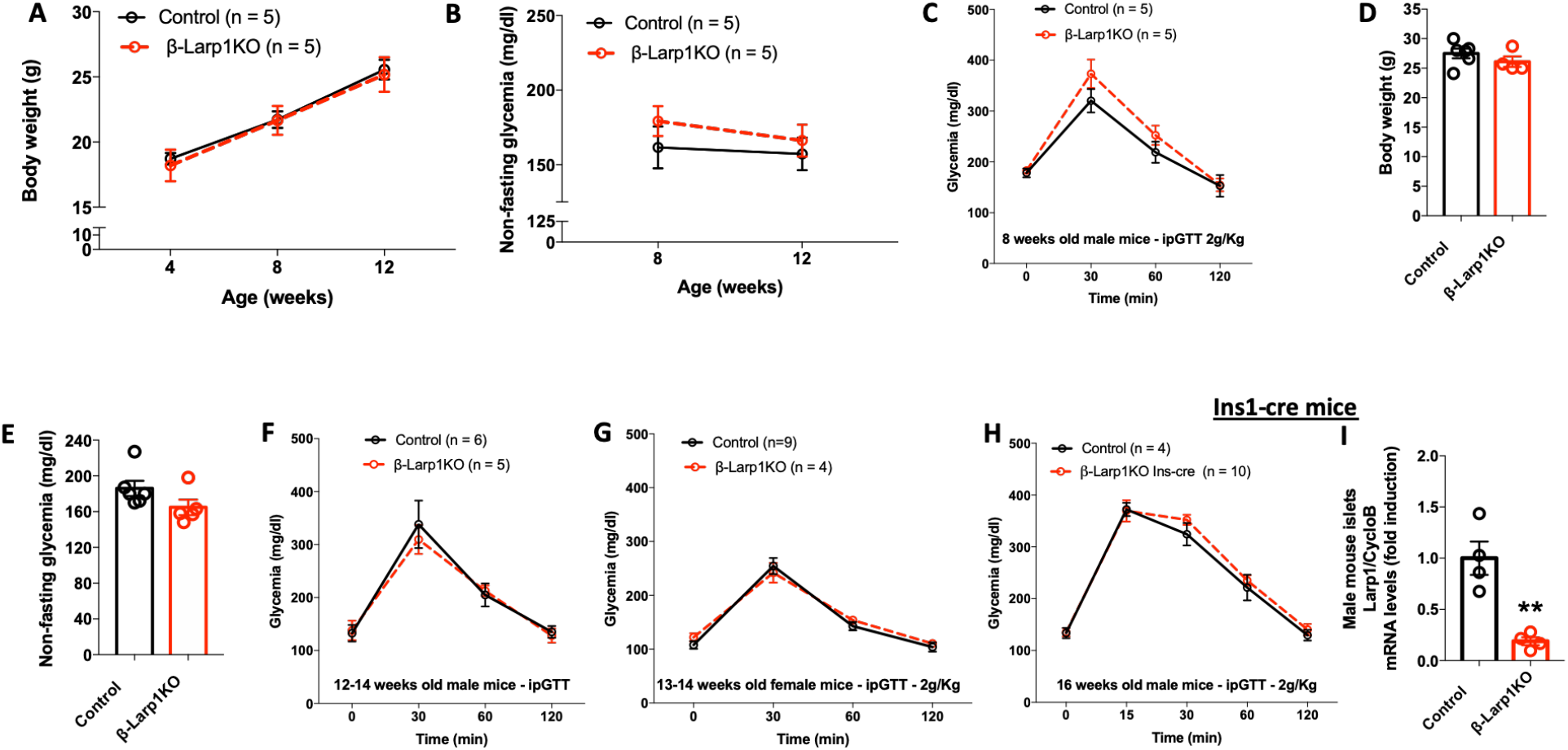
Males and females β-Larp1KO mice have normal glucose homeostasis. **(A)** Body weight gain, **(B)** fed glycemia and **(C)** intraperitoneal glucose tolerance test (ipGTT) in the first cohort of male *β-Larp1KO* mice; **(D)** Body weight, **(E)** fed glycemia and **(F)** ipGTT in the second cohort of male *β-Larp1KO* mice; **(G)** ipGTT in female *β-Larp1KO* mice and **(H)** ipGTT in male *β-Larp1KO* mice generated by crossing the *floxed-Larp1* mice with the *Ins1-cre* mice instead of *Rip-Cre* used in A-G. ** p < 0.01 compared to control mice assessed by Student’s T test.

**Figure 5.**
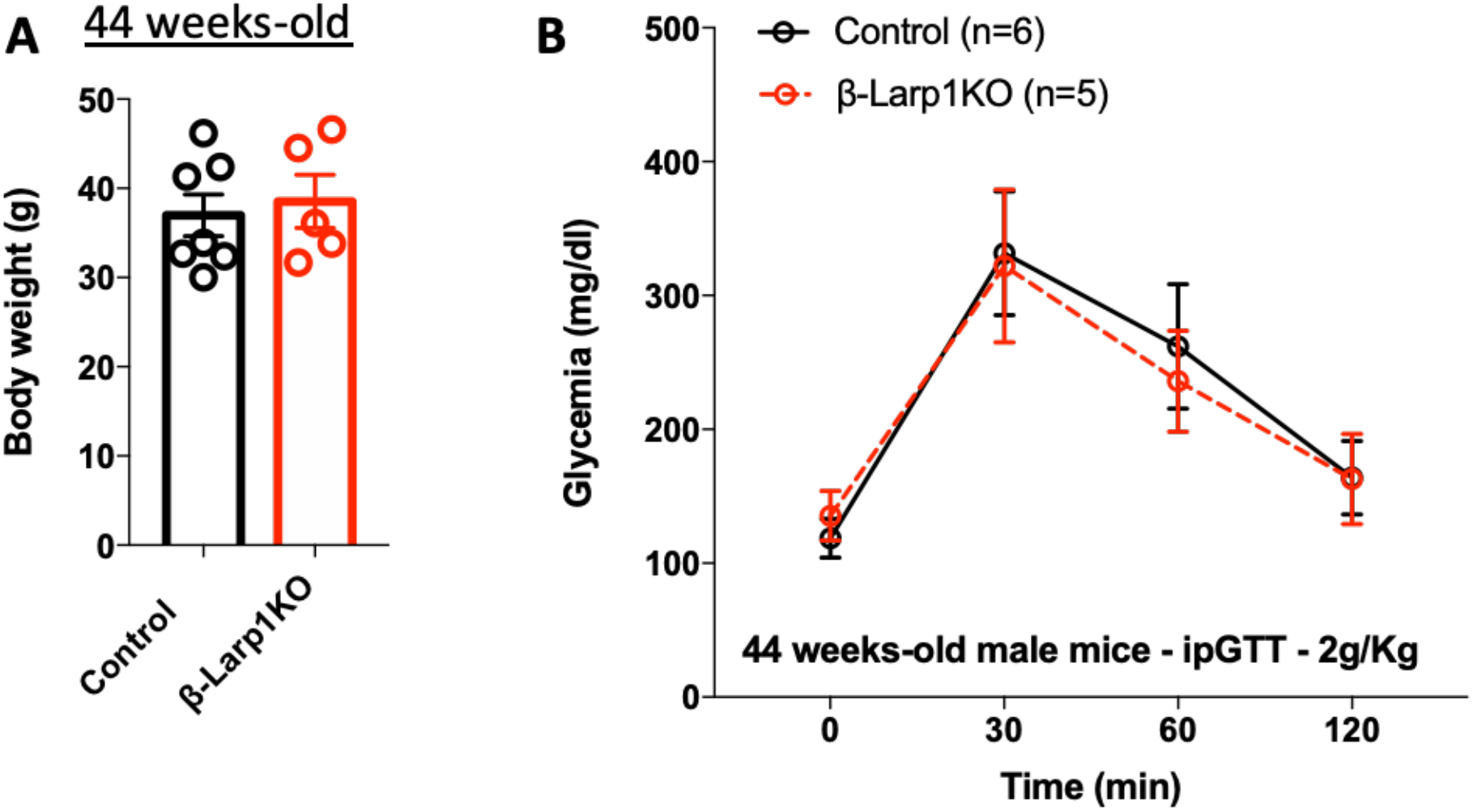
Aged β-Larp1KO mice are tolerant to glucose as control mice. **(A)** Body weight and **(B)** intraperitoneal glucose tolerance test in male *β-Larp1KO* mice at 44 weeks of age.

### Exposure of β-Larp1KO mice to high fat diet did not alter glucose homeostasis

It is well-characterized that high fat diet (HFD) provokes insulin resistance, increasing the demand for insulin production and secretion by β-cells, resulting in higher insulinemia. Insulin in turn stimulates β-cell expansion through the activation of Akt/mTORC1 pathway. Therefore, we decided to challenge the *β-Larp1KO* mice under HFD. We placed the first cohort used in Fig 4A under HFD. The *β-Larp1KO* mice gained weigh as much as control mice (Fig 6A and B). There was no difference in fed glycemia before, during and after 8 weeks under HFD (Fig 6C). Intraperitoneal glucose tolerance test at 4 and 8 weeks after HFD was comparable between *β-Larp1KO* and control mice (Fig 6D and E). We also tested whether incretins could play a role in the *β-Larp1KO* mice by performing an oral glucose tolerance test and found no difference between the groups (Fig 6F).

**Figure 6.**
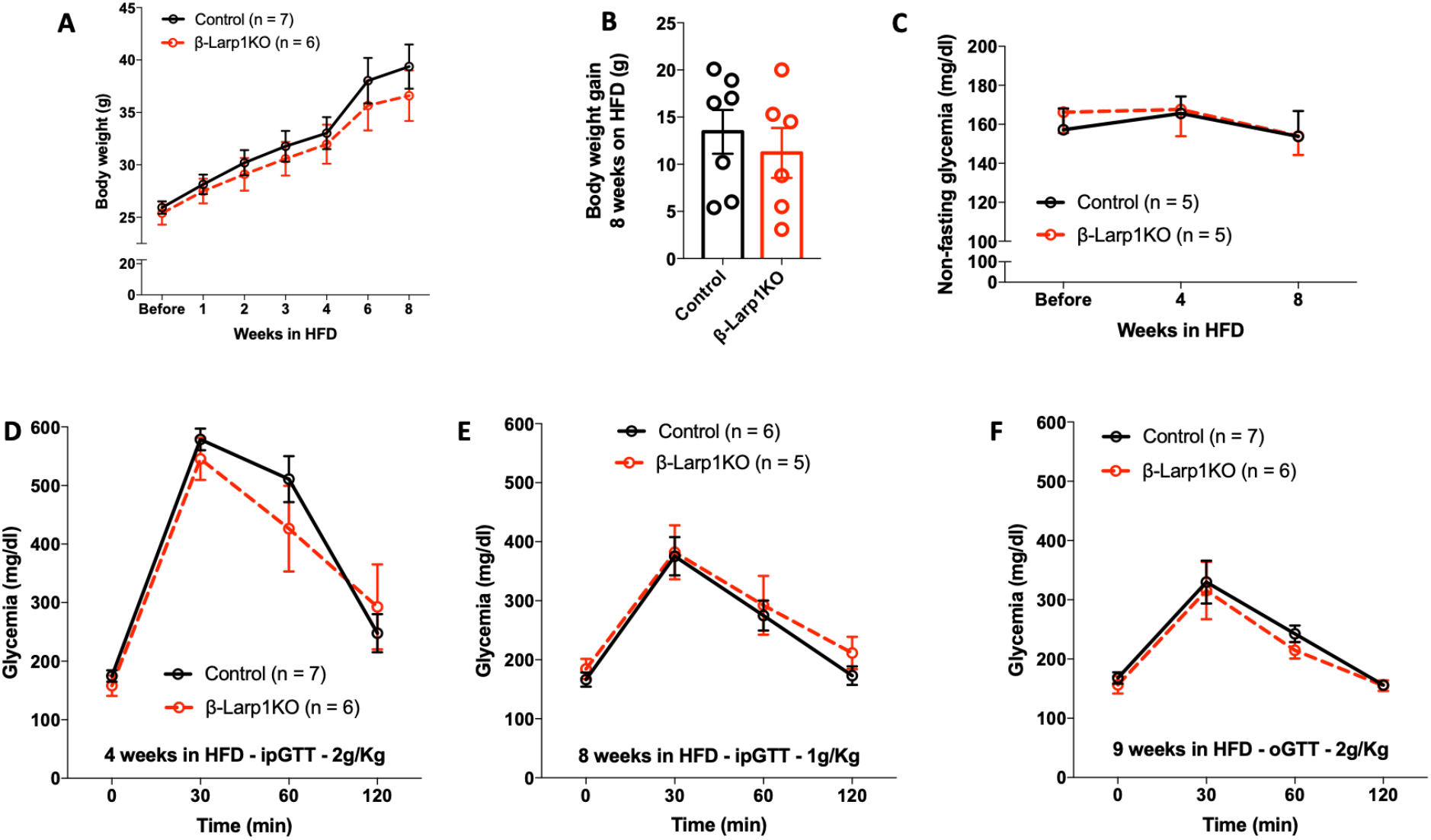
β-Larp1KO mice under high fat diet. **(A)** Body weight, **(B)** Body weight gain and **(C)** fed glycemia of mice fed with high fat diet (60% fat) for 8 weeks. Intraperitoneal glucose tolerance test (ipGTT) after **(D)** 4 and **(E)** 8 weeks in HFD. **(G)** Oral glucose tolerance test (ipGTT) after 9 weeks in HFD.

### Long-term exposure to high branched-chain amino acid diet slightly impairs glucose tolerance in β-Larp1KO mice

As an alternative method, we placed the second-cohort used in Fig 4D and E under branched-chain amino acids diet (BCAA). BCAA diet directly stimulate mTORC1 activity, especially the enriched L-leucine amino acid (Condon and Sabatini 2019). There was no difference in body weight gain in the *β-Larp1KO* and control groups (Fig 7A and B). Glucose tolerance was the same between the groups at 4 and 8 weeks after high BCAA diet (Fig 7C and D). However, after 16 weeks, *β-Larp1KO* were slightly intolerant to glucose compared to littermate control mice (Fig 7E and F). Oral tolerance to glucose was the same after 17 weeks in BCAA diet (Fig 7G).

**Figure 7.**
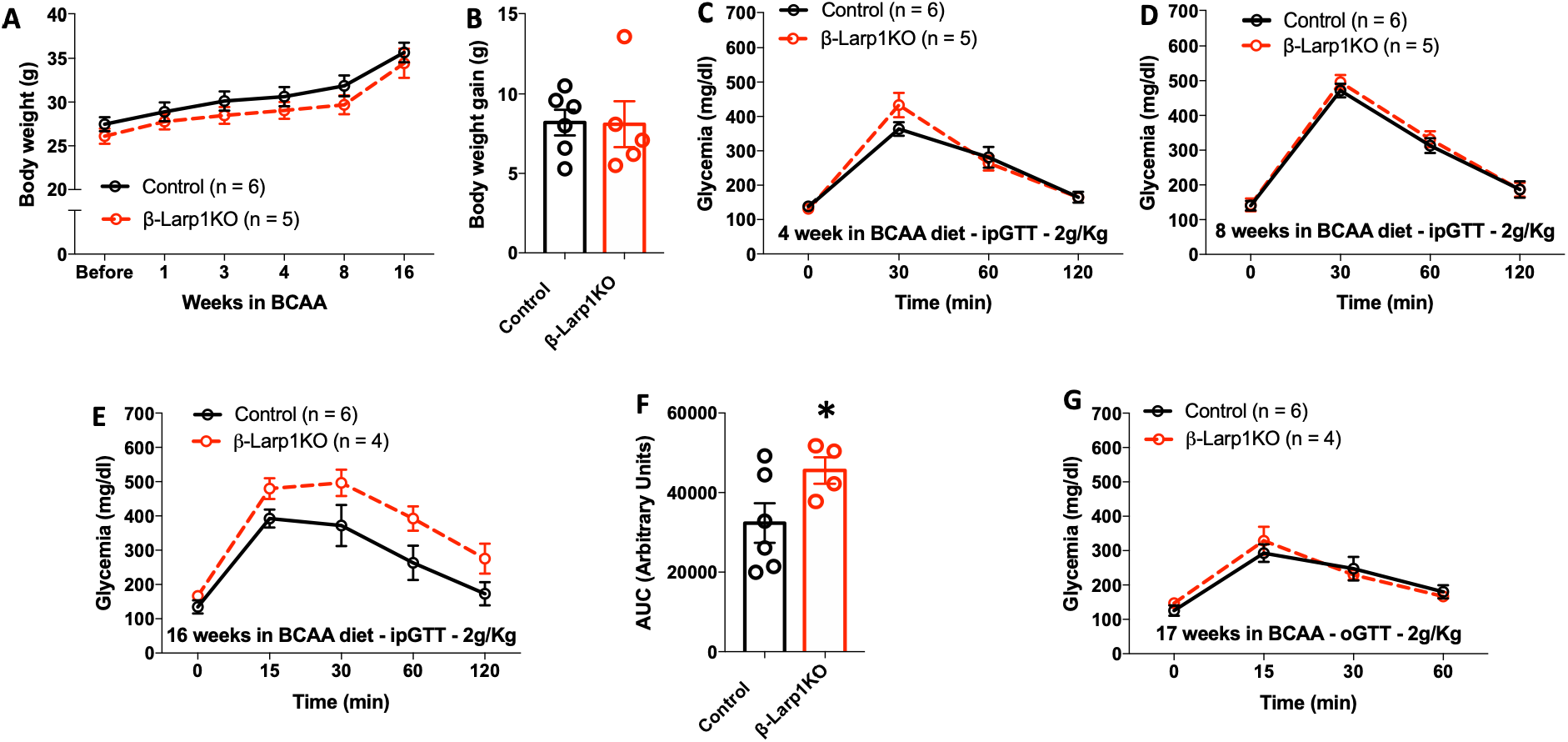
β-Larp1KO mice under high branched-chain amino acid diet (BCAA). **(A)** Body weight and **(B)** Body weight gain of mice fed with high branched-chain aminoacid diet (150% BCAA) for 16 weeks. **(C)** Intraperitoneal glucose tolerance test (ipGTT) after 4 **(D)** 8 and **(E)** 16 weeks in BCAA diet. **(F)** Quantification of the area under the curve in the ipGTT performed in **E**. **(G)** Oral glucose tolerance test (ipGTT) after 17 weeks in BCAA diet. * p < 0.05 compared to control mice assessed by Student’s T test.

## DISCUSSION

We report herein that La related protein 1 (LARP1) is highly expressed in human β-cells and mouse islets compared to the other members of the family. Furthermore, type 2 diabetes up-regulates LARP1 and LARP1B consistent with high protein translation, cell proliferation and mTORC1 activity (Yuan, Rafizadeh et al. 2017, Ardestani, Lupse et al. 2018). However, mice lacking LARP1 specifically in β-cells (*β-Larp1KO* mice) developed normally and glucose metabolism was similar to control mice. Even under diet-induced stress (high fat or diet high in branched-chain amino acid), glucose metabolism in *β-Larp1KO* mice did not deviate from control mice. There was only a minor impaired glucose tolerance after long-term (16 weeks) in BCAA diet. Therefore, we conclude that LARP1 is dispensable for pancreatic β-cell function and glucose homeostasis.

A constant cellular regulation of protein synthesis and breakdown determine cellular function and growth. During the progression of diabetes, β-cells expand in size and number to meet the high metabolic demand imposed by insulin resistance (Alejandro, Gregg et al. 2015). mTORC1 signaling pathway is highly activated in β-cells from diabetic patients and rodents, indicating enhanced protein synthesis and increased proliferation (Yuan, Rafizadeh et al. 2017). The current studies were designed to interrogate the *in vivo* role of LARP1 in β-cell function based on previous finding that LARP1 and mTORC1 work together to regulate the translation of key mRNAs, the 5’ TOP mRNA (Hong, Freeberg et al. 2017, Lahr, Fonseca et al. 2017, Philippe, Vasseur et al. 2018). This group of mRNA encodes components of the protein translation machinery. We found that LARP1 is the most expressed LARP in mouse islets and human β-cells. Moreover, LARP1 is up-regulated in β-cells in patients diagnosed with type 2 diabetes. This is consistent with the *in vitro* findings that LARP1 regulates proliferation (Tcherkezian, Cargnello et al. 2014, Hong, Freeberg et al. 2017, Berman, Thoreen et al. 2020). In several different types of cancer, highly phosphorylated LARP1 has been reported, although kinases other than mTORC1 are likely to induce LARP1 phosphorylation in cancer (Hopkins, Mura et al. 2016, Ye, Lin et al. 2016, Xu, Xu et al. 2017, Berman, Thoreen et al. 2020).

Surprisingly, we found that *β-Larp1KO* mice exhibit a normal phenotype. mTORC1 plays a fundamental role in β-cell physiology by controlling 5’ cap-dependent translation of critical proteins in β-cell (Blandino-Rosano, Barbaresso et al. 2017, Ni, Gu et al. 2017). mTORC1 activity in β-cells is higher during embryological development and in the first week of post-natal maturation, followed by lower activity levels in mature β-cells (Ni, Gu et al. 2017, Sinagoga, Stone et al. 2017, Jaafar, Tran et al. 2019, Helman, Cangelosi et al. 2020, Katsumoto and Grapin-Botton 2020). mTORC1 is reactivated in diabetic states and the chronic hyperactivation could play a role in β-cells dysfunction or failure (Yuan, Rafizadeh et al. 2017, Ardestani, Lupse et al. 2018). The fact that mTORC1 and LARP1 interact to each other to control cellular protein translation capacity, and, more importantly, that mTORC1 activity and LARP1 expression are both increased in diabetes prompted us to generate a conditional mouse strain to disrupt LARP1 specifically in β-cells. The reasons for the normal glucose metabolism in the *β-Larp1KO* mice are not clear. We observed an 80% reduction of *Larp1* gene expression in islets of *β-Larp1KO* mice. This is similar to the 80% reduction in mTORC1 signaling found in the *β-RaptorKO* mice generated by crossing the *floxed-raptor* mouse with the same *RIP-Cre* mouse used to produce the *β-Larp1KO* mice (Blandino-Rosano, Barbaresso et al. 2017). The RIP-Cre-induced recombination is ~90-95% of all insulin positive cells (Alejandro, Bozadjieva et al. 2017, Blandino-Rosano, Barbaresso et al. 2017). We speculate that the remaining 20% expression in mouse islets is coming from non-β-cells, very few cells scaping from recombination and minor acinar contamination of the islet preparation. Due to ectopic cre-recombinase expression outside the pancreas in the *RIP-cre* mice, we confirmed the neutral metabolic phenotype in the *β-Larp1KO* mice by using the *Ins1-Cre* mouse. Therefore, it is unlikely that normal β-cell function in the *β-Larp1KO* mice is due to poor cre-mediated recombination.

An alternative explanation to the lack of phenotype in the *β-Larp1KO* mice would be an upregulation of other members of the family. The LARP family consists of six members: LARP1, 2 (1B), 4, 5 (4B), 6, and 7 (Bousquet-Antonelli and Deragon 2009, Hong, Freeberg et al. 2017). They all contain the RNA-binding domain but only LARP1 and LARP1B present the DM15 motif and interact with mTORC1 (Bousquet-Antonelli and Deragon 2009, Hong, Freeberg et al. 2017). In the *β-Larp1KO* mice, islet LARP1B expression was similar to control littermates. Although there was trend to higher levels of LARP6 in *β-Larp1KO* mice, this is unlikely to explain the normal phenotype as LARP6 is barely expressed in mouse islets (< 5%) and even a 3-fold induction would still result in very low levels of LARP6. However, it is possible that expression of other LARP family members at normal or slightly increase levels (LARP6) is sufficient to maintain protein translation and β-cell function.

The small increase in glucose levels observed in the intraperitoneal glucose tolerance test in the *β-Larp1KO* mice undergoing long-term high BCAA diet (16 weeks) opens the possibility that the lack of LARP1 potentially limits protein synthesis in prolonged and sustained mTORC1 activation as in diabetogenic conditions. However, the upregulation of LARP1 in diabetes (Fig 1B) might attenuate the absence of LARP1 in the responses to HFD or BCAA diet.

In summary, LARP1 is highly expressed in human β-cells and mouse islets, and is upregulated in diabetes. However, LARP1 is dispensable for pancreatic β-cell function and glucose homeostasis *in vivo*. β-cell adaptation to chronic exposure to high concentration of branched-chain amino acids may require LARP1.

## ACKNOWLEDGEMENTS

The authors declare no conflict of interest. We would like to appreciate the technical support provided by Portia Ritter and Pau Romaguera Llacer. The current studies were funded by the National Institute of Health (NIDDK) grant R01-DK073716 and DK084236.

